## Supplementary figures and images for "CASK and FARP localize two classes of post-synaptic ACh receptors thereby promoting cholinergic transmission"

### Figure S1, related to figure 2, lin-7 and lin-10.tif

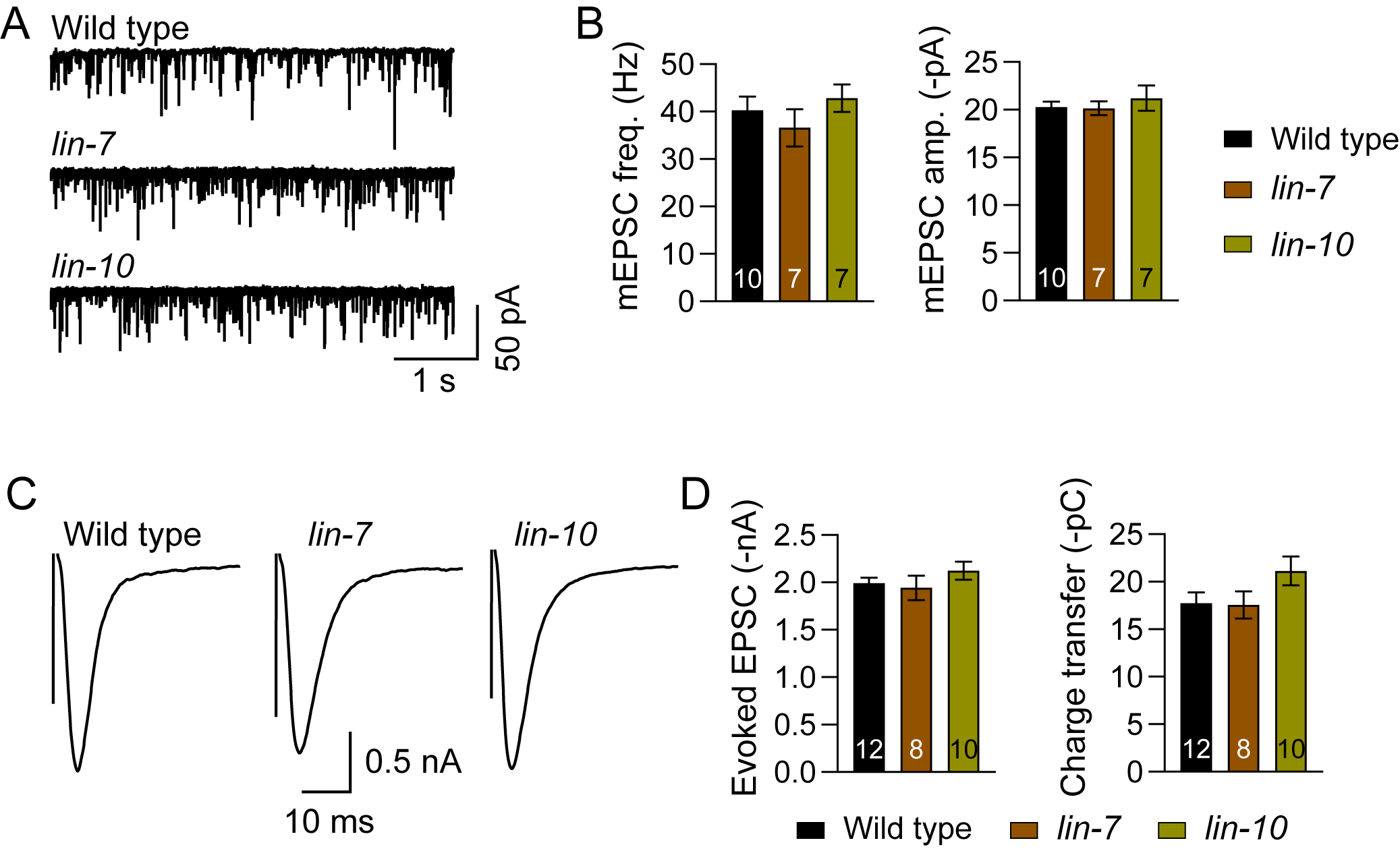

### Figure S2, related to figure 2, acr-16.tif

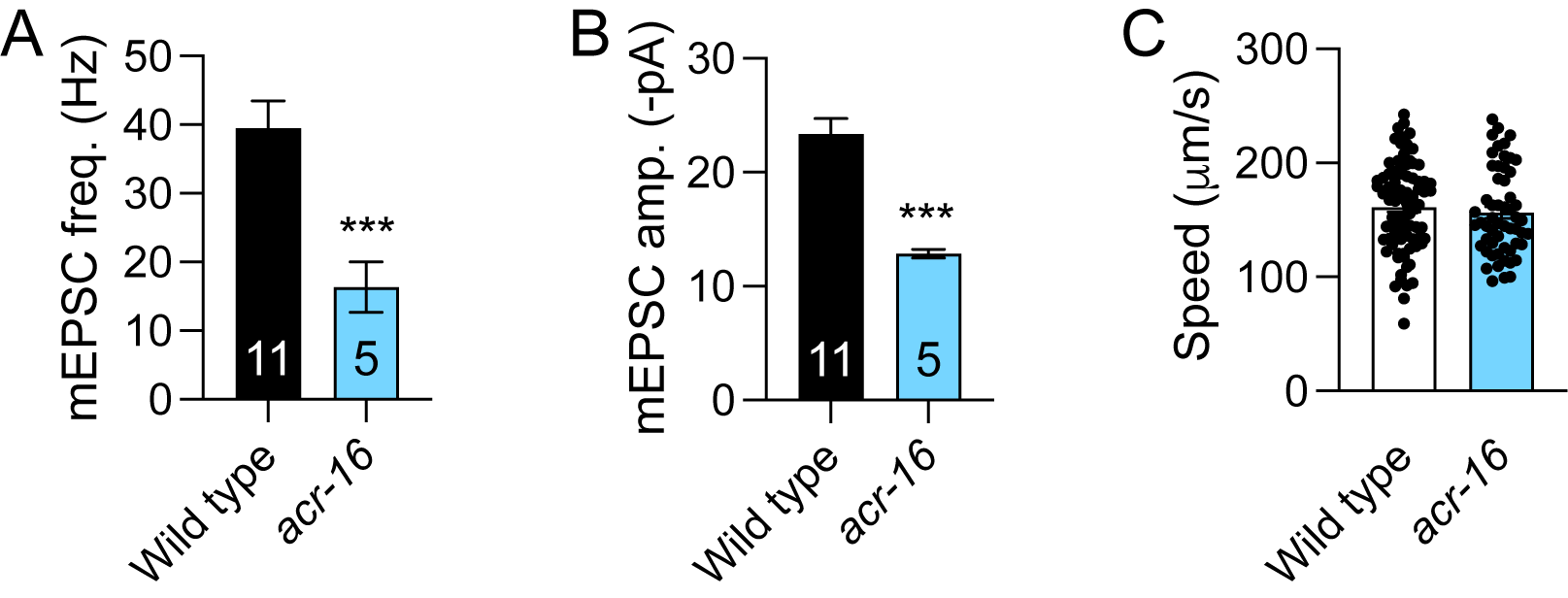

### Figure S3, related to figure 3, cKO speed.tif

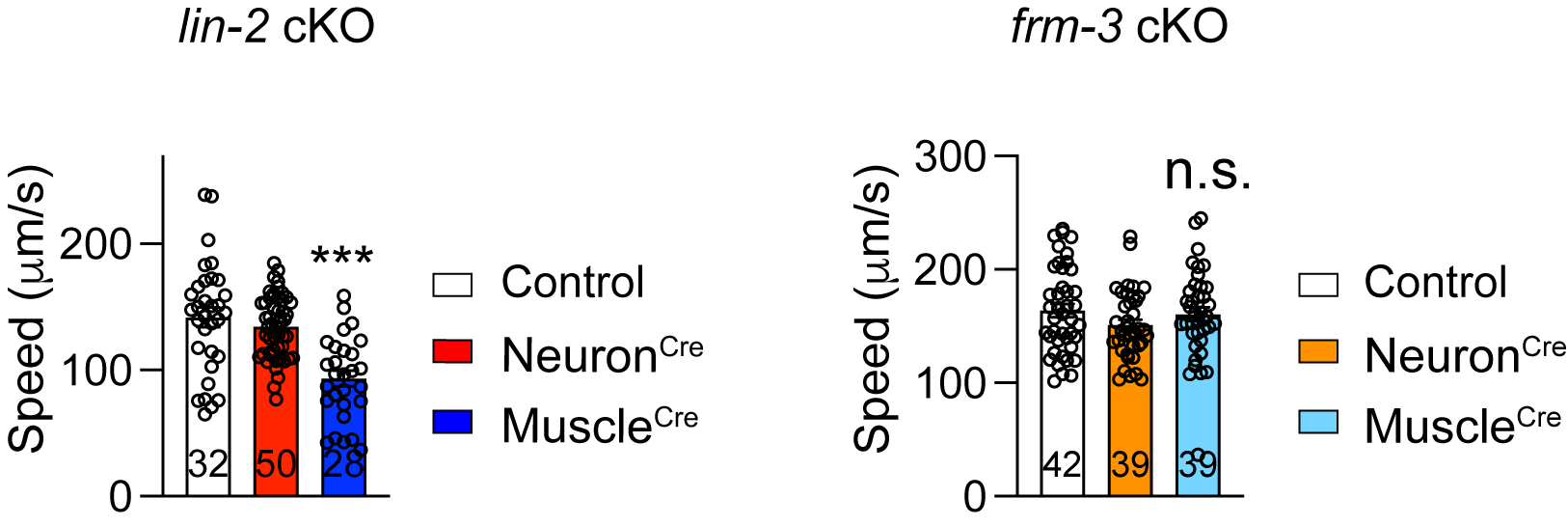

### Figure S4, related to figure 3, UNC-2 Cre from Josh.tif

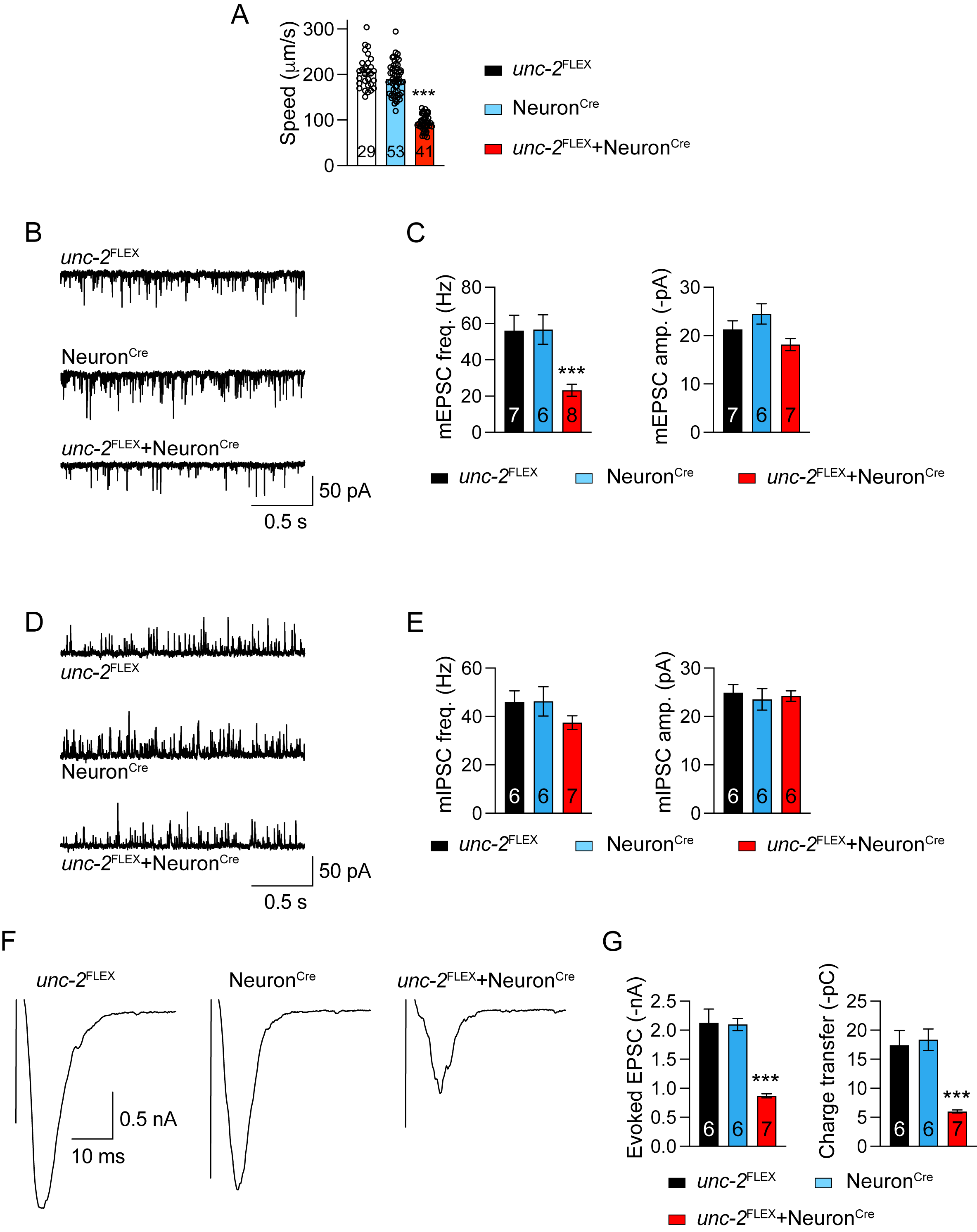

### Figure S5, related to figure 4, mini decay.tif

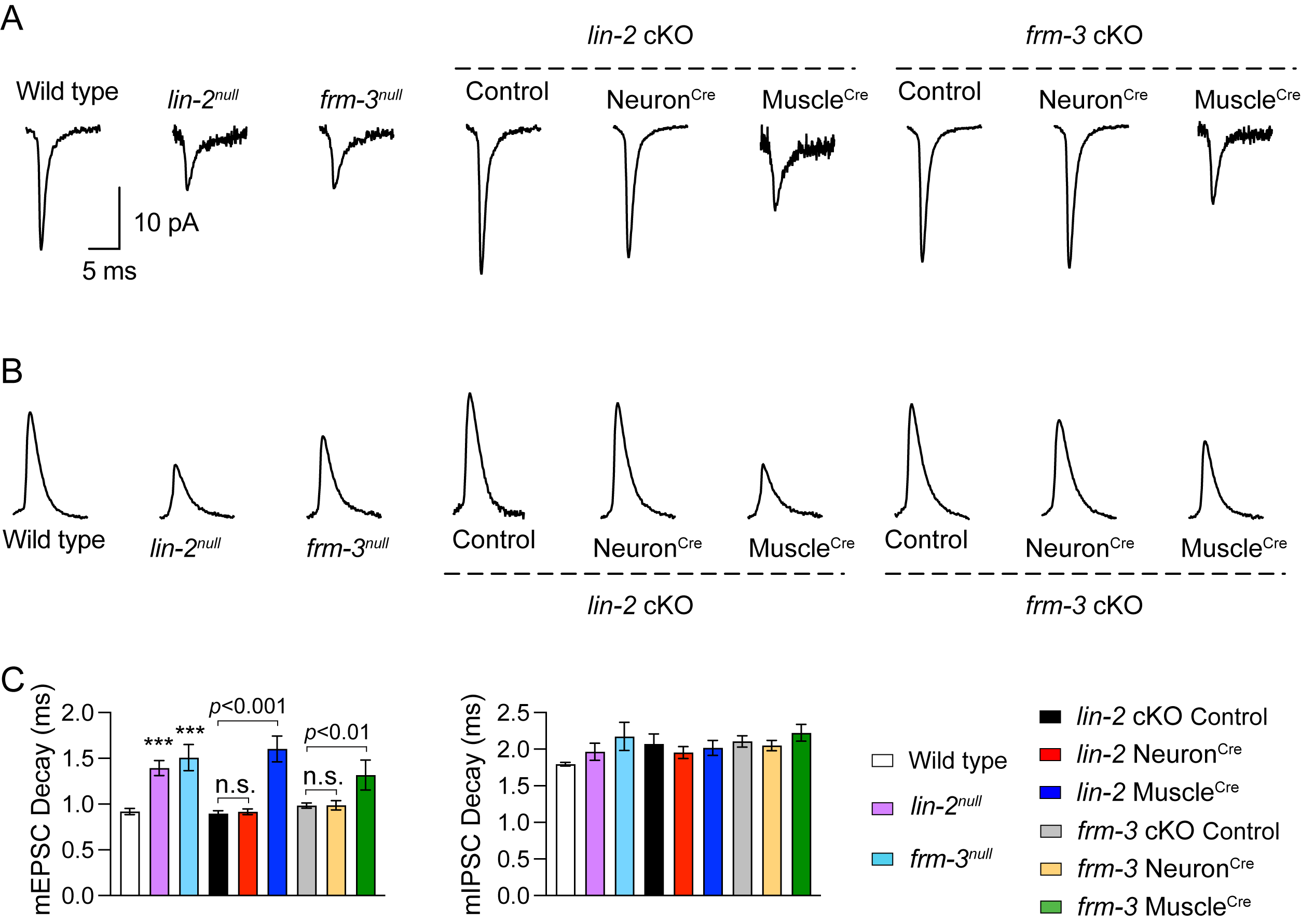

### Figure S6, related to figure 4, cKO puff.tif

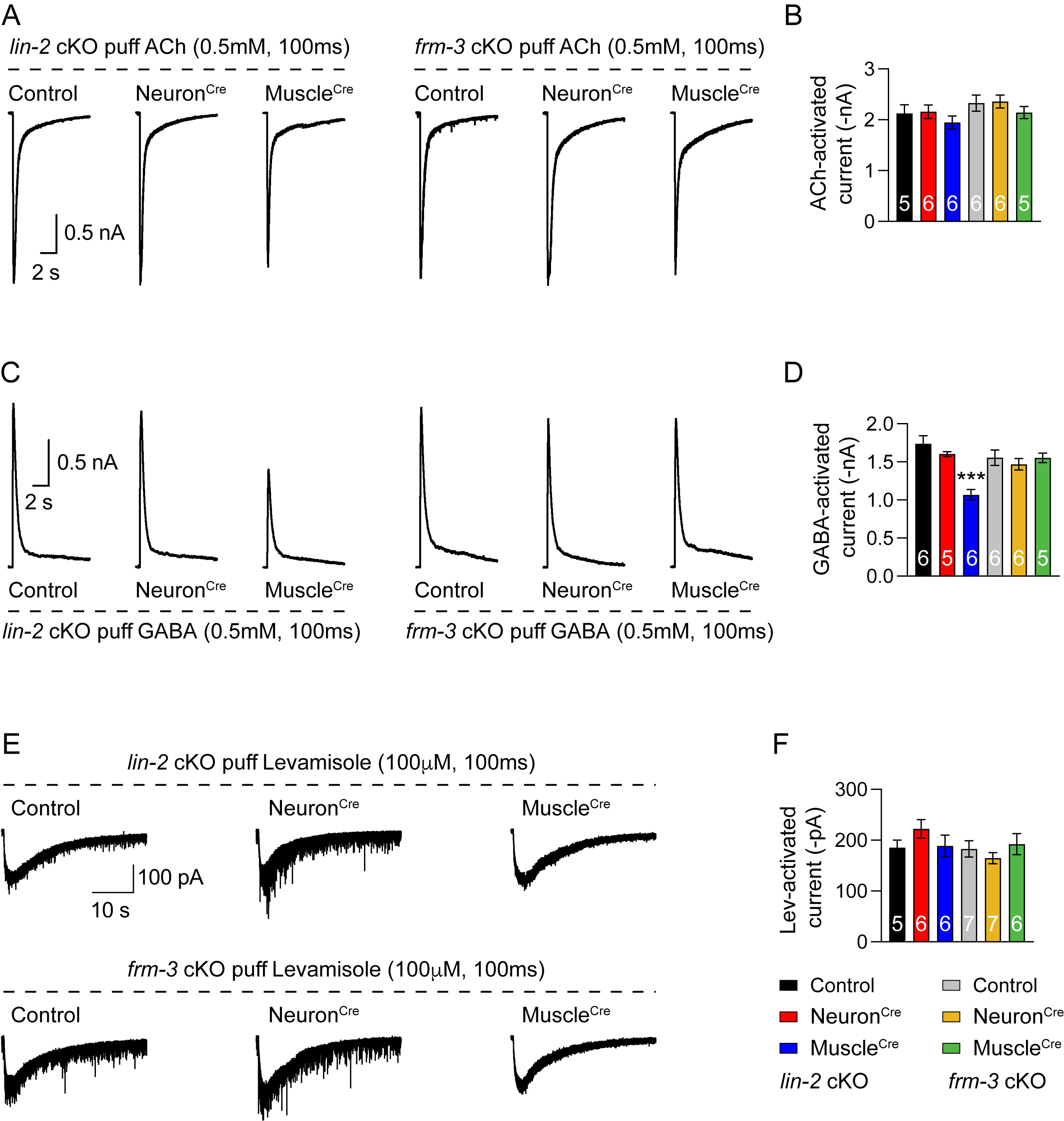
